# Deep-sea plastisphere: long-term colonization by plastic-associated bacterial and archaeal communities in the Southwest Atlantic Ocean

**DOI:** 10.1101/2021.01.26.428295

**Authors:** Luana Agostini, Julio Cezar Fornazier Moreira, Amanda Gonçalves Bendia, Maria Carolina Pezzo Kmit, Linda Gwen Waters, Marina Ferreira Mourão Santana, Paulo Yukio Gomes Sumida, Alexander Turra, Vivian Helena Pellizari

## Abstract

Marine plastic pollution is a global concern because of continuous release into the oceans over the last several decades. Although recent studies have made efforts to characterize the so-called plastisphere, or microbial community inhabiting plastic substrates, it is not clear whether the plastisphere is defined as a core community or as a random attachment of microbial cells. Likewise, little is known about the influence of the deep-sea environment on the plastisphere. In our experimental study, we evaluated the microbial colonization on polypropylene pellets and two types of plastic bags: regular high density polyethylene (HDPE) and HDPE with the oxo-biodegradable additive BDA. Gravel was used as control. Samples were deployed at three sites at 3,300 m depth in the Southwest Atlantic Ocean and left for microbial colonization for 719 days. For microbial communities analysis, DNA was extracted from the biofilm on plastic and gravel substrates, and then the 16S rRNA was sequenced through the Illumina Miseq platform. Cultivation was performed to isolate strains from the plastic and gravel substrates. Substrate type strongly influenced the microbial composition and structure, while no difference between sites was detected. Although several taxa were shared among plastics, we observed some groups specific for each plastic substrate. These communities comprised taxa previously reported from both epipelagic zones and deep-sea benthic ecosystems. The core microbiome (microbial taxa shared by all plastic substrates) was exclusively composed by low abundance taxa, with some members well-described in the plastisphere and with known plastic-degradation capabilities. Additionally, we obtained bacterial strains that have been previously reported inhabiting plastic substrates and/or degrading hydrocarbon compounds, which corroborates our metabarcoding data and suggests the presence of microbial members potentially active and involved with degradation of these plastics in the deep sea.

## 1. Introduction

Plastic waste has become a global challenge. Although many countries are involved in public policies to mitigate this problem, tons of plastic waste continue to enter the oceans annually (GESAMP, 2019; Jambeck et al., 2015; PlasticEurope, 2019; Schmidt et al., 2017). As a consequence, marine plastic contamination is ubiquitous, including in remote marine environments such as the deep ocean (Woodal et. al., 2014). In the marine environment, plastic waste is exposed to physical and chemical factors that contribute to the degradation and fragmentation of this material into microplastics (i.e. plastic objects < 5 mm in diameter) (Roager and Sonnenschein, 2019). Microplastic chemical composition, density and shape determine whether it is more likely to float or sink, which will influence the distribution in pelagic or benthic ecosystems (Cole et al., 2011; Pierdomenico et al., 2019; Pinnell and Turner, 2019; Van Cauwenberghe et al., 2013; Woodall et al., 2014). Although both macro- and microplastics have already been reported in deep-ocean regions, knowledge regarding plastic colonization by deep-sea prokaryotic communities in both pelagic and benthic ecosystems is incipient (Bergmann and Klages, 2012; Chiba et al., 2018; Galgani et al., 1996; Krause et al., 2020; Pierdomenico et al., 2019; Schlining et al., 2013; Woodall et al., 2018).

Plastics can be used as substrata and be rapidly colonized by microorganisms, which form biofilms on the plastic surface, a community we refer to as the “plastisphere” (Zettler et al., 2013). The ability to colonize and metabolize substrate surfaces is a mechanism that promotes advantages for microorganisms in situations with nutritional limitation (Dang and Lovell, 2016), as can be the case in deep-sea regions. Plastic substrates in the deep ocean may offer a new hotspot of colonization, as well as a relevant source of carbon to support the microbial community, proportionally magnifying the abundance of potentially plastic degrading microorganisms within the plastisphere. A growing number of studies have reported microorganisms capable of degrading hydrocarbons (Didier et al., 2017; Oberbeckmann et al., 2017), raising the hypothesis that they would be consuming this material in nutritionally limited conditions, such as those found in deep-sea ecosystems.

In addition to the influence of the environment (which can be prone to microbial colonization or not), polymer composition is suggested to modulate the structure and composition of the plastisphere (Dussud et al., 2018a; Dussud et al., 2018b; Kirstein et al., 2018, 2019; McCormick et al., 2014; Pinto et al., 2019; Zettler et al., 2013). Moreover, geographic locations have also been indicated to exert influence on the microbial community of the plastisphere (Oberbeckmann et al., 2016). In contrast, independent of environmental factors or plastic substrate type, some microbial taxa have been reported in multiple plastispheres (Tu et al., 2020; Zettler et al., 2013) and they are now referred to as “core microbiome”. The core microbiome members comprise taxa with high occupancy across a dataset that are hypothesized to reflect functional relationships with the host (or substrate) (Shade and Handelsman, 2012). In the plastisphere, these members are thought to be involved in biofilm formation and/or metabolizing compounds from the plastic substrate (Didier et al., 2017; Tu et al., 2020). Bacterial families such as Oleiphilaceae and Hyphomonadaceae have been found colonizing multiple substrates (Bryant et al., 2016; De Tender et al., 2017; Oberbeckmann et al., 2018; Pinto et al., 2019; Zettler et al., 2013). The known capacity of these families to metabolize hydrocarbon compounds may be the key functional trait enabling them to colonize different plastics (Golyshin et al., 2002; Ozaki et al., 2007).

In the last decade, the number of studies on the plastisphere has increased (Amaral-Zettler et al., 2020). However, efforts have been concentrated mainly on epipelagic (e.g. Bryant et al., 2016; Carson et al., 2013; Zettler et al., 2013) or shallow benthic systems (Pinnel and Turner, 2019). Only a few studies, however, analyzed the plastisphere in deep-sea habitats (Krause et. al., 2020; Woodal et. al, 2018). Studies that assessed marine plastisphere microbial communities can be classified into three basic groups: those that randomly collected plastic marine debris (PMD) or microplastics (Bryant et al., 2016; De Tender et al., 2015; Zettler et al., 2013); studies that deployed plastic substrates in the ocean (Oberbeckmann et al., 2016; Tu et al., 2020) and those conducted in laboratory conditions (Kirstein et al., 2018; Ogonowski et al., 2018). The microbial community structure inhabiting deployed plastic substrates for long periods is as yet poorly studied (Kirstein et al., 2018; Oberbeckmann et al., 2016, 2018; Tu et al., 2020). Based on these studies, the marine plastic bacterial community is mainly represented by the Gammaproteobacteria, Alphaproteobacteria and Bacteroidetes classes, while the archaeal community is represented by the Euryarchaeota and Crenarchaeota phyla (Eriksen et al., 2014; Oberbeckmann et al., 2018; Oberbeckmann and Labrenz, 2020; Quero and Luna, 2017; Urbanek et al., 2018; Woodall et al., 2018).

Here, we conducted the first *in situ* experimental study that characterized the structure and composition of the microbial community (the bacterial and archaeal) associated with different types of plastic substrates deployed for 719 days at three sites in deep waters in the Southwest (SW) Atlantic Ocean (3,300 m). Our objectives in this study were to understand (i) if there are differences in microbial communities inhabiting plastics polymers as opposed to control samples and adjacent seawater, and differences among multiple sites, (ii) if there is a core microbiome among plastics that can contribute to the description of the plastisphere in the deep SW Atlantic Ocean (different from shallow waters), and (iii) if it is possible to isolate viable bacteria through cultivation of plastic substrates that are potentially related to plastic degradation. For this, we performed a high-throughput sequencing of the 16S rRNA gene to access bacterial and archaeal communities and used traditional culturing methods to assess the plastics as substrates for microbial growth.

## 2. Materials and Methods

### 2.1. Studied area and deployment methods

Experimental sites are located in a region that encompasses the transition between the North Atlantic Deep Water (NADW) and the Lower Circumpolar Deep Water (LCDW). This region is characterized by water temperature between 3 °C and 4 °C, salinity between 34.6 and 35.0, oxygen concentration above 5 mL L^−1^ and low nutrient levels (i.e. oligotrophic) (Gonzalez-Silvera et al., 2004; Krueger et al., 2015).

Autonomous aluminum structures that housed multiple experiments, called landers, were deployed at three sites along the southeastern Brazilian continental margin, and between 21°S and 28°S. The sites were named ES (22°50’27.24”S; 38°24’58.68”W), RJ (25°20’17.88”S; 39°38’28.32”W) and SP (28°1’42.24”S; 43°31’43.32”W), with distances of 304 km between ES and RJ, 786 km between ES and SP, and 496 km between RJ and SP (Figure 1). Landers were deployed in June 2-6, 2013 at 3,300 m depth, using the R/V Alpha Crucis from the Oceanographic Institute at Universidade de São Paulo (IO-USP). On each lander, single use plastic bags of two different polymers, 60g of pristine PP pellets (Braskem) and 60 g of inorganic material as a control were deployed. Plastic bags included: (i) regular high density polyethylene (HDPE) grocery bag material (Valbags, ValGroup Brasil), (ii) biodegradable grocery bag material made from HDPE with the oxo-biodegradable additive BDA (Willow Ridge Plastics, Inc.). All types of substrates were placed inside fiberglass mesh bags (~8 × 15 cm; mesh size, 1 mm), attached to the metal lander frame and secured with nylon fishing line and rope before deployment.

**Figure 1:**
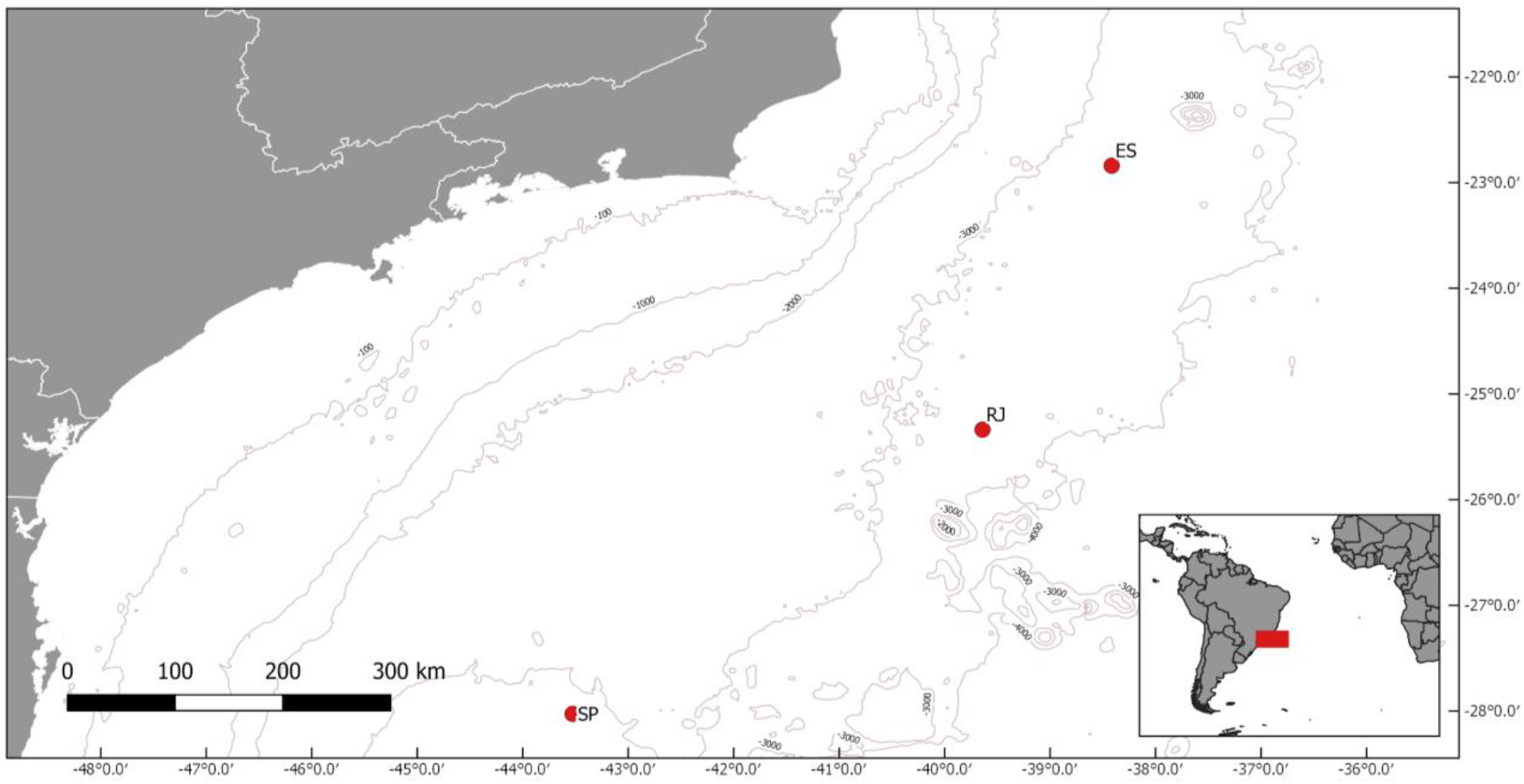
Map of the study region (Southwestern Atlantic) indicating the three experimental sites where the landers were deployed (red dots), all placed along the 3,300 meter bathymetric line. Sites: ES, Espírito Santo; RJ, Rio de Janeiro; SP, São Paulo. Source: GeoMapApp^®^.

After 719 days in all cases, 23.6 months (on May 22-25, 2015), samples were collected with the support of NPo Almirante Maximiano (H-41, Brazilian Navy). For DNA extraction, 30 pellets of gravel, plastic and cut strips of plastic bag material were rinsed lightly with autoclaved distilled water. These were then placed into vials (10 per vial for pellets or sufficient to occupy approximately the same vial volume as pellets for bags, n = 3 replicate vials), filled with RNAlater buffer solution (Thermo Fisher Scientific, Waltham, MA, USA), and stored at −20 °C until analysis. For live culturing, an additional 5 pellets or cut strips of plastic bag material were placed directly into Eppendorf tubes and filled with sterilized seawater without rinsing. Eppendorf tubes were shaken gently, and stored at 4 °C until analysis. All handling materials were sterilized between every step.

Before retrieving each lander, water samples from the same depth and current as the plastic samples were collected using a Rosette water sampler equipped with Niskin bottles. Water samples (adjacent water - AW) were collected to analyze the structure of the microbial communities in the environment where plastics were deployed. Each water sample collected (~ 10 L) was immediately filtered through a 0.22 μm polycarbonate membrane (diameter 45 mm; Millipore, Bedford, MA, USA) using a peristaltic pump, and stored at −80 °C.

### 2.2. DNA extraction, 16S rRNA gene amplification and sequencing

Samples were processed at LECOM, the Microbial Ecology Laboratory at the Oceanographic Institute (IO-USP) of University of São Paulo. Extraction of the total DNA from the plastics was performed in triplicate using the PowerSoil DNA Isolation Kit (MoBio, Carlsbad, CA, USA). Extraction of DNA from the water was performed using a PowerWater DNA Isolation Kit (MoBio, Carlsbad, CA, USA) according to the manufacturer’s specifications.

Six PCR reactions from each sample were pooled and purified with the DNA Clean & Concentrator™ kit (Zymo Research, Irvine, CA, USA), and quantified with Qubit 1.0 fluorometer (Thermo Fisher Scientific, Waltham, MA, USA) and the Qubit^®^ dsDNA HS Assay Kit (Thermo Fisher Scientific, Waltham, MA, USA). PCR was conducted following the Earth Microbiome 16S Illumina Amplicon Protocol. Amplicons were sent to the Molecular Research - MR. DNA company (Texas, USA) for sequencing on the Illumina Miseq platform in a 2×300 bp pair-end system. The V4 hypervariable region of the 16S rRNA gene from Bacteria and Archaea was amplified with the universal primers 515F and 806R (Caporaso et al., 2010) with specific adapters for the Illumina Miseq platform. Sequence data (raw .fastq files) have been submitted to the GenBank under accession number PRJNA692207.

### 2.3. Bioinformatics and statistical analyses

Processing and quality control of reads was performed using QIIME2 version 2019.10 (Bolyen et al., 2019). After graphic inspection of quality profiles, raw reads were subjected to trimming and filtering, then clustered into Amplicon Sequence Variants (ASV) with DADA2 denoising (Callahan et al., 2016) using the QIIME 2 package (Bolyen et al., 2019). Sequence counts were rarefied to 45,020 reads per sample across all samples to mitigate uneven sequencing depth.

The ASV richness, Shannon and InvSimpson diversity indexes were calculated using *phyloseq* and *vegan* packages. Normality and Homogeneity of variances was assessed by Shapiro-Wilk normality and Levene’s test, respectively. If the data showed a normal distribution and the variances were homogeneous, differences between groups were assessed by one-way analysis of variance (ANOVA) and subsequent post-hoc Tukey’s tests, which were performed using *stats* and *agricolae* packages in R (v.3.6.1) to assess differences in diversity indexes among substrates and sites. To compare the structure of the bacterial communities among substrates and sites, non-metric multidimensional scaling (NMDS) ordination was performed, based on weighted unifrac dissimilarities among all samples. Differences in the microbial community structure among substrates and sites were tested by performing a permutational multivariate analysis of variance (PERMANOVA) on the community matrix (Anderson, 2001).

To identify an ASV that was significantly more abundant among substrate types, we performed DESeq2, Differential Expression analysis for Sequence count data (Love et al., 2014). The DESeq2 input was a rarefied microbial dataset previously treated using the Prevalence Interval for Microbiome Evaluation (PIME) package (Roesch et al., 2020). PIME uses machine-learning to generate ASV prevalence among samples, and validate it by comparison with control Monte Carlo simulations with randomized variations of sequences (Roesch et al., 2020). The full rarefied dataset consisting of 5,199 ASVs was filtered using the PIME R package (Roesch et al., 2020). PIME removes the within-group variations and captures only biologically significant differences which have high sample prevalence levels. PIME employs a supervised machine-learning algorithm to predict random forests and estimates out-of-bag (OOB) errors for each ASV prevalence dataset at 5% intervals. High OOB errors indicate that a given prevalence dataset interval is noisy, while the minimal OOB errors (OBB error = zero) represent the absence of noise. Here, the minimal OOB errors occurred with a 70% prevalence interval, which represented 471,078 sequences distributed among 535 ASVs. This 70% prevalence dataset was used for DESeq2 subsequent analyses.

To observe the occurrence of ASVs among substrate types, the samples were grouped by substrate type and the taxa abundance table transformed to presence/absence. The unique and shared ASVs were then visualized using an UpSet plot, *UpSetR* package (Conway et al., 2017). The ASVs shared by all plastic types were considered the core microbiome.

### 2.4. Cultivable Plastic-associated Bacteria

In sterile Petri dishes, samples of plastic substrates were inoculated directly into the mineral culture medium described by Sekiguchi et al., (2010). The medium was prepared to contain per liter of distilled water: 1.87% of Marine Broth (Difco), 1.5% NaCl, 0.35% KCl, 5.4% MgCl_2_. 6H_2_O, 2.7% MgSO_4_. 7H_2_O, 0.5% CaCl_2_. 2H_2_O, 1.2% agar with 0.25% poly-β-hydroxybutyrate (PHB) granules added. The samples were incubated for 15 days at 13 °C, or until the growth of colonies around the plastic samples was observed. All morphologically different macroscopic colonies were selected using the depletion technique two to three times until pure colonies for sequencing were obtained. The isolates were preserved in 20% glycerol in an ultra-freezer at −80 °C.

The genomic DNA of 22 isolates was extracted using the Purelink Genomic DNA kit (Invitrogen by Thermo Fisher Scientific, Carlsbad, USA), according to the manufacturer’s specifications. Amplification of the RNAr 16S gene was conducted using primers 515F (5 ’- GTGYCAGCMGCCGCGGTAA - 3’) and 1401R (5 ’- CGGTGTGTACAAGGCCCGGGA - 3’). The polymerase chain reaction (25 μL reaction) was performed using Gotaq Mix Hot Start, 0.25 μL of each primer and 2 μL of DNA template. The PCR conditions were: initial denaturation temperature of 95 °C, 3 min; followed by 30 cycles of 94 °C, 1 min; 53 °C, 30 seconds; 72 °C, 1 min; and a final extension at 72 °C for 10 min. The PCR product was purified using the DNA Clean & Concentrator kit (Zymo Research, Irvine, USA) according to the manufacturer’s specifications and sent for sequencing at Genomic Engenharia Molecular, where they were sequenced using the BigDye Terminator v3.1 Cycle Sequencing Kit (Applied Biosystems, Palo Alto, USA) with the 515F primer.

Sequence analysis was initially performed using CodonCodeAligner Software (CodonCode Corporation, Dedham, MA, USA). Through this software, the sequences were checked for quality and treated. After obtaining the treated sequences, the SILVA v138 database (High-Quality Ribosomal RNA Databases) was used to align the sequences, to identify the isolates and to construct the phylogenetic trees through MEGA X software (Kumar et al., 2018), using the maximum-likelihood method (999 bootstraps). All sequencing data was deposited in GenBank (National Center for Biotechnology Information Sequence Read Archives) under accession numbers between MW216888 and MW216902.

## 3. Results

### 3.1. ASV richness and Alpha diversity among substrata and sites

From the 15 samples sequenced, a total of 1,086,988 valid sequences (i.e. reads) were obtained, representing an average of 72,465±12,068 (SD) reads per sample. The obtained reads were clustered into 5,310 ASVs, representing an average of 702±149 ASVs per sample. Rarefaction curves indicated a stationary phase, suggesting sufficient depth of sequencing to account for the diversity of the microbial community on the plastic substrates, gravel, and seawater samples (Figure S1).

Overall, among substrate type the ASV richness measured by the Chao1 estimator was significantly higher on gravel samples (898.33±84.77) (ANOVA; F = 4.85; df = 4; p = 0.027), while the microbial richness from AW samples and plastic substrates (HDPE and PP) was lower (631±106, 616±80 and 583±141, respectively) (Figure S2A). Although the Shannon diversity was not affected by substrate types (ANOVA; F = 2.77; df = 4; p = 0.102), the diversity observed on gravel samples was 1.05, 1.07, and 1.2-fold higher than PP, HDPE-OXO, and HDPE samples, respectively (Figure S2B). The evenness was also not affected significantly by substrate type (ANOVA; F = 1.65; df = 4; p = 0.111), but a lower mean evenness was observed on HDPE (0.66), AW (0.68), and HDPE-OXO (0.71), while the PP samples showed the higher evenness (0.76) (Figure S2C).

The site did not exhibit a significant effect on any diversity indexes measured. However, the ASV richness measured by the Chao1 estimator was higher from RJ samples (741.09±159.06) (ANOVA; F = 0.61; df = 2; p = 0.564), while SP and ES showed similar richness (675.54±81.16; 681.38±203.29, respectively) (Figure S2D). Shannon index averages were high in RJ samples (4.78±0.55) and SP (4.71±0.20), while ES showed the lowest mean values (4.51±0.55) (ANOVA; F = 0.38; df = 2; p = 0.517) (Figure S2E). Finally, the lowest evenness was observed in the ES samples (0.69±0.07) and similar between RJ and SP (0.72±0.06 and 0.72±0.02, respectively) (ANOVA; F = 0.61; df = 2; p = 0.565) (Figure S2F).

### 3.2. Microbial community structure among substrata and sites

Non-metric multidimensional scaling (NMDS) based on weighted Unifrac dissimilarities revealed that the global pattern of microbial diversity was significantly explained by the substrate type (Figure 2). Based on Permutational Multivariate Analysis of Variance (PERMANOVA) the microbial community structure was highly dependent on the substrate type (R^2^ = 0.78, P = 0.001), while the site effect was lower and not significant (R^2^ = 0.11, P = 0.054). Adjacent water (AW) samples were particularly distinct, while gravel samples showed more similarities to plastic samples.

**Figure 2.**
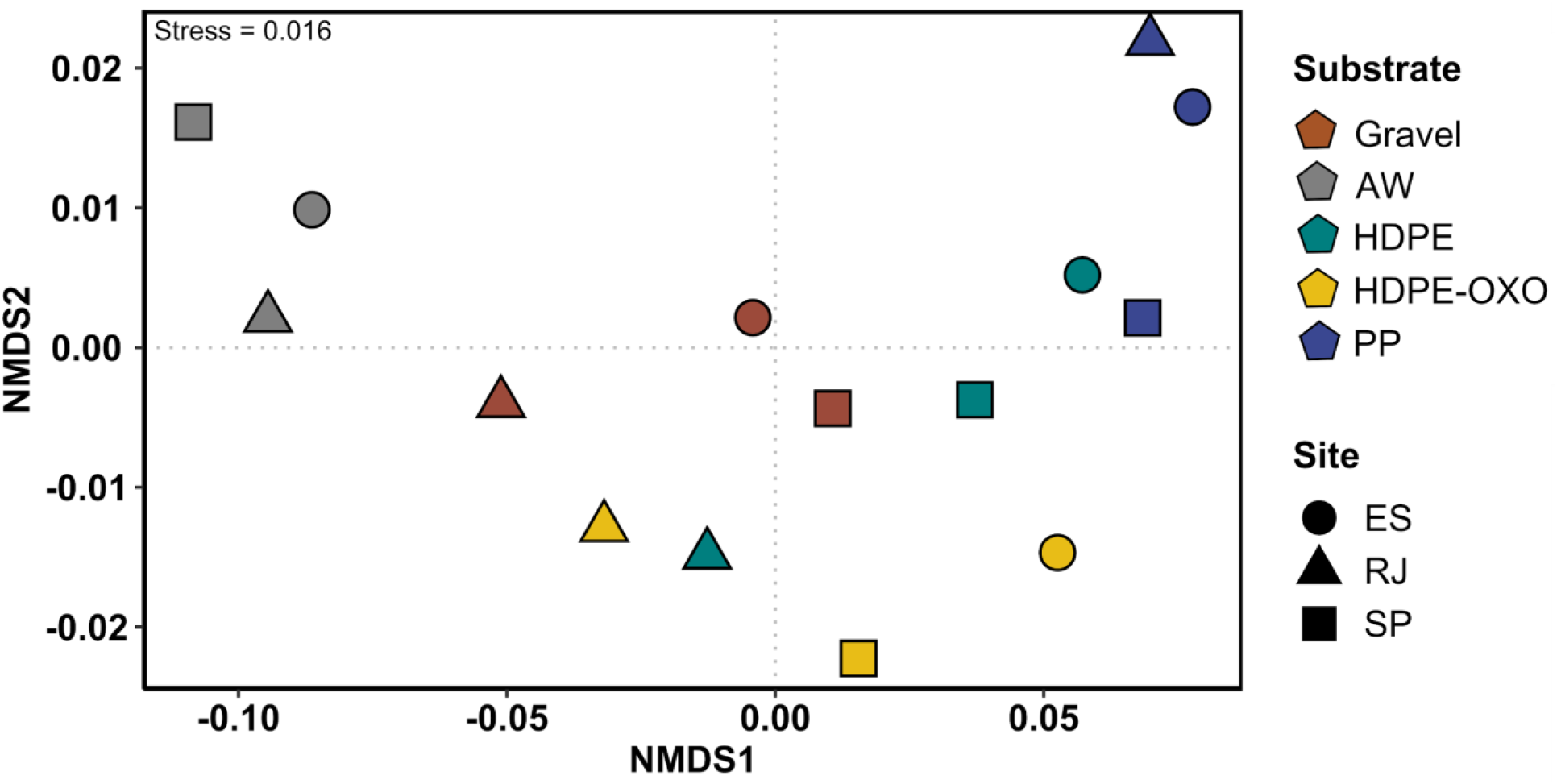
Non-metric multidimensional scaling (NMDS) ordination of weighted unifrac dissimilarities in microbial community structures. Substrate: Gravel; AW, adjacent water; HDPE, High density polyethylene bag (film); HDPE-OXO, High density polyethylene bag (film) with oxo-biodegradable additive BDA; PP, Polypropylene pellets. Sites: ES, Espírito Santo; RJ, Rio de Janeiro; SP, São Paulo.

### 3.3. Microbial composition and taxa differential abundance between plastic and control samples

The microbial community of all substrates were composed in majority by the phylum Proteobacteria (40 to 77%), while other phyla showed different colonization patterns amongst plastic substrates (HDPE, HDPE-OXO and PP), gravel and AW, including both presence/absence and differences in relative proportions of the phyla (Figure 3A). For example, Firmicutes was abundant in all PP samples (average 20±6%), while found in smaller proportions in HDPE (6.6±6.2%), HDPE-OXO (6.1±5.6%) and gravel samples (3.8±1.9%). Among plastics, NB1-j was more abundant in HDPE (4.6±5.9%) and archaeal phyla (such as Crenarchaeota and Nanoarchaeota) were prevalent in HDPE and HDPE-OXO (5.1±4 and 7.3±4.7%, respectively). The lowest proportions of those archaeal phyla were observed in PP (0.7±0.4%). Chloroflexi, Marinimicrobia (SAR406 clade) and SAR324 (Marine group B) were mainly present in AW samples (7.1±1.6, 5.6±0.28 and 6.9±0.1%, respectively) but not in the plastic substrates used in this study.

**Figure 3.**
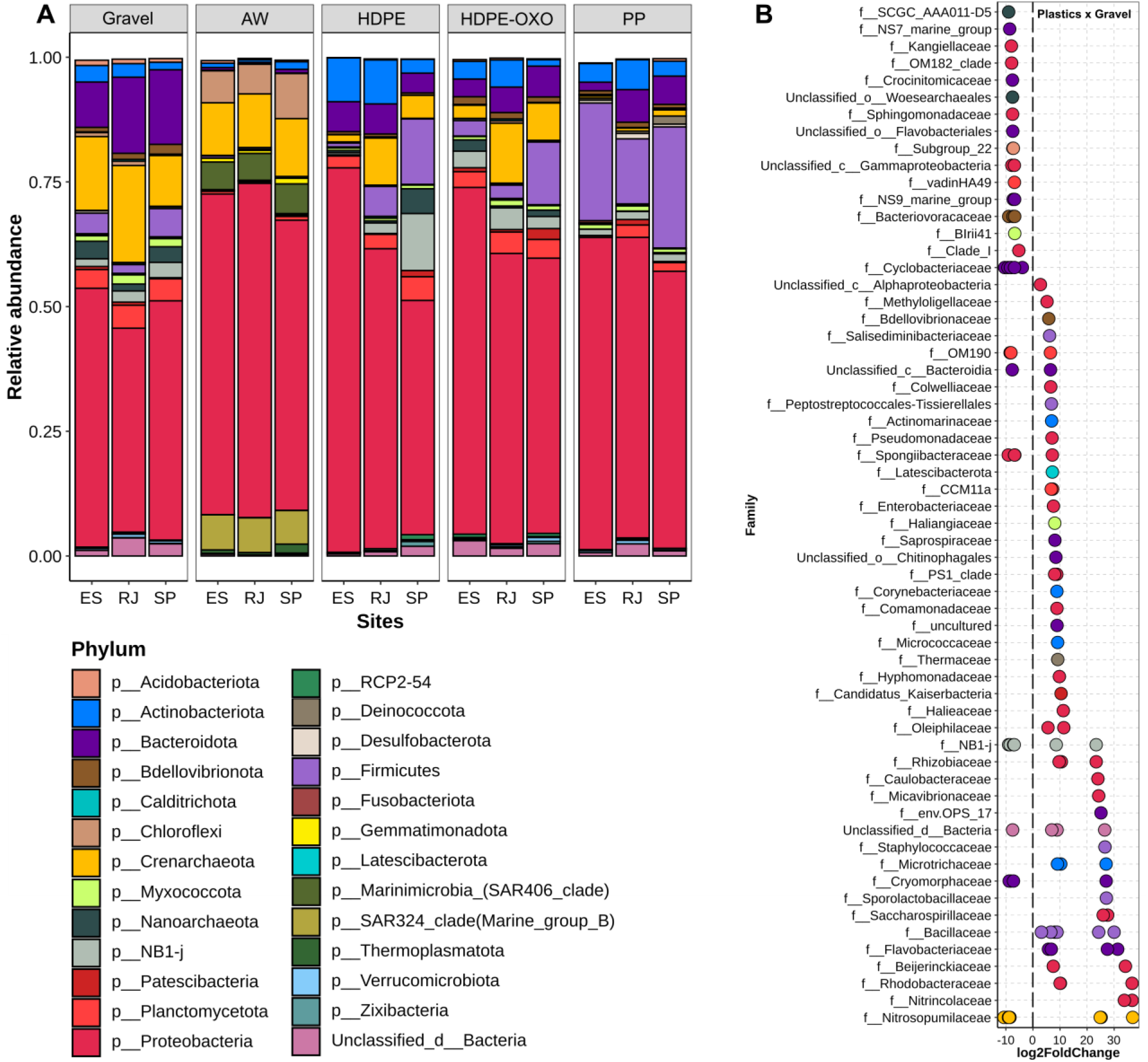
(A) Relative abundance of the microbial community at the Phylum level (Bacteria and Archaea) among sites. (B) Differentially abundant Amplicon Sequence Variant (ASV) comparing plastic substrates (HDPE, HDPE-OXO and PP merged) and gravel. Substrate: Gravel; AW, adjacent water; HDPE, High density polyethylene; HDPE-OXO, High density polyethylene with oxo-biodegradable additive BDA; PP, Polypropylene. Significant ASVs (padj < 0.05) are represented by single data points, grouped by family on the y-axis and by color according to the taxonomic phylum from which the ASV originates. Positive values (log2FoldChange) indicate ASVs significantly more abundant in plastic substrates; Negative values indicate the opposite. Unclassified taxons were represented by the prefix Unassigned_.

Differences between plastics and gravel samples were strongly evident when we examined deeper taxonomic levels. We observed ASVs from a total of 37 families that were significantly more abundant in plastic substrates when compared with gravel samples; these include families, such as Methylologellaceae, Colwelliaceae, Pseudomonadaceae, Haliangiaceae, Micrococcaceae, Halieceaea, Oleiphilaceae, Flavobacteriaceae, Rhizobiaceae, Microtrichaeae, Rhodobacteraceae, among others (Figure 2B).Seven families showed ASVs with differential abundances for both plastic and gravel samples, such as Cyclobacteriaceae, OM190, Spongiibacteraceae, NB1-j, Cryomorphaceae, Nitrosopumilaceae and unclassified Bacteroidia. Gravel samples exhibited 16 families significantly more abundant in comparison with plastics (e.g. Kangiellaceae, OM182 clade, Crocinitomicaceae, Sphingomonadaceae and Bacteriovoracaceae) (Figure 2B).

### 3.4. Microbial composition and taxa differential abundance among plastic types

Among each substrate type, the largest number of unique ASVs was found in gravel samples (1096 ASVs), followed by AW, PP, HDPE-OXO and HDPE, with respective values of 954, 736, 732 and 475 ASVs. Grouping those ASVs into families to examine abundance, we identified a predominance of low abundance families on all substrates deployed (relative abundance < 2% of total community) on PP (71.9%), HDPE-OXO (63.9%), gravel (56.8%) and HDPE (40.2%). The percentage of low abundance families in AW samples was lower (21.4%) (Figure 4).

**Figure 4.**
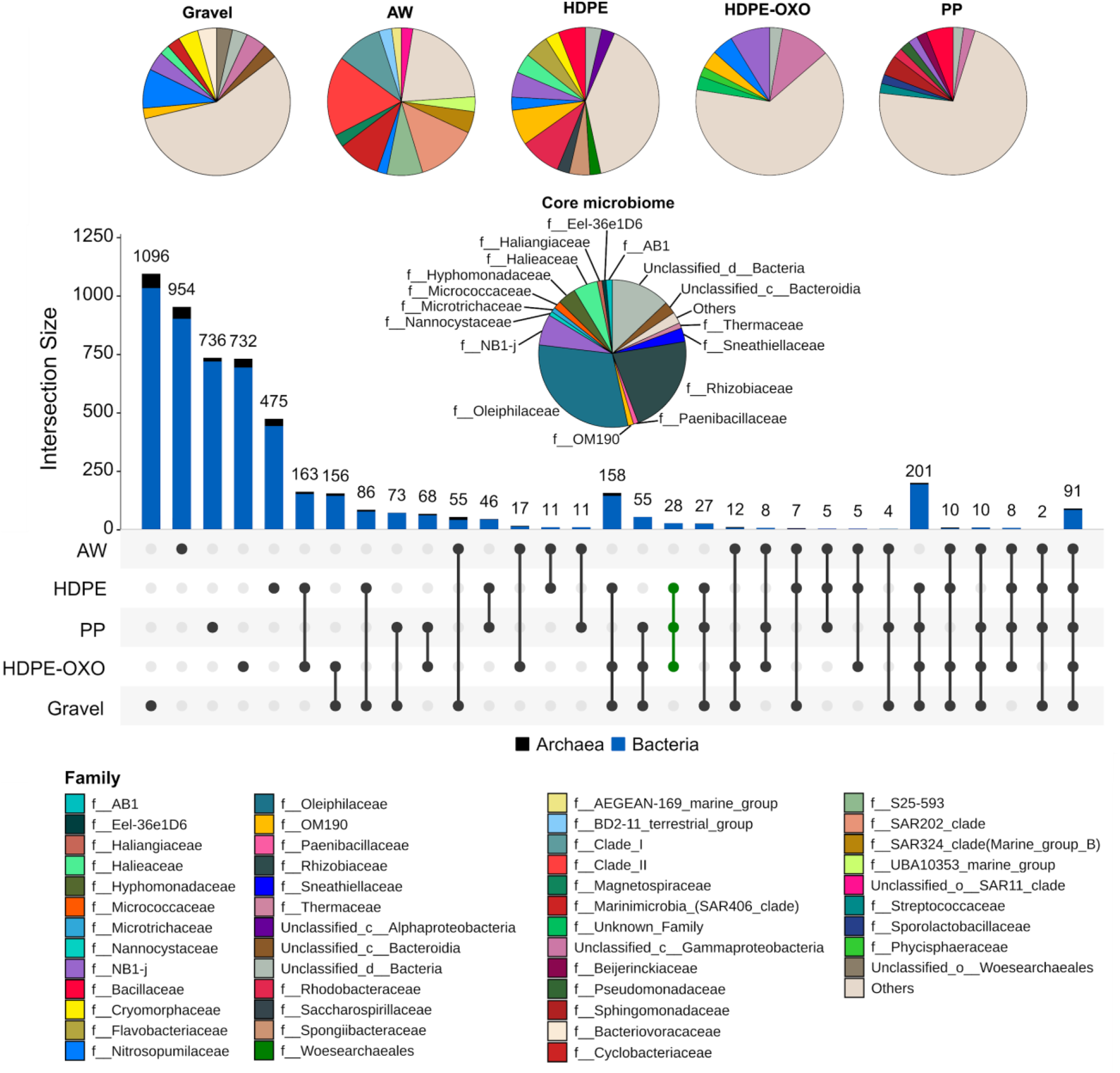
Upset plot composed by ASVs identified among substrates. Circles indicate substrates. Black lines connecting circles indicate shared ASVs. Vertical bars indicate intersection size (number of ASVs) on each set. Blue and black bars represent Bacteria and Archaea ASVs, respectively. The green line represents ASVs shared by all plastic samples. Substrates: AW, adjacent water; HDPE, High density polyethylene; HDPE-OXO, High density polyethylene with biodegradable additive BDA; PP, Polypropylene. Pie charts show microbial composition specific to each substrate (families with abundance > 2%) and those shared among plastics types (core microbiome). Substrate: Gravel; AW, adjacent water; HDPE, High density polyethylene; HDPE-OXO, High density polyethylene with oxo-biodegradable additive BDA; PP, Polypropylene.

Overall, the number of common ASVs amongst substrates deployed (pairwise combinations among HDPE, HDPE-OXO, PP and gravel) was higher than common ASVs between AW and substrates deployed (Figure 4). These results may confirm taxa effectively colonizing the substrates over long periods. Another piece of evidence to support this idea is the high number of ASVs (201) shared among all substrates deployed, while only 91 ASVs were observed shared among the substrates deployed and the adjacent water (AW) (Figure 4).

Similar polymer composition was also suggested as exerting influence on the microbial taxa composition. We identified 163 ASVs shared between HDPE and HDPE-OXO, plastic substrates composed basically by HDPE. In contrast, a lower number of ASVs were shared between PP and HDPE-OXO (68) and PP and HDPE (46), substrates composed of different polymers (Figure 4).

Based on the 44 families with ASVs significantly more abundant in plastic substrates than in gravel samples (Figure 2B), we checked the distributions of these families amongst plastic substrates types (Figure 5). Results showed us three major family groups: (i) the generalists, found with significant abundance in all plastic types (HDPE, HDPE-OXO, and PP), (ii) the plastic HDPE generalists, found with significant abundance in plastic HDPE substrates (HDPE and HDPE-OXO), and finally, (iii) the specialists group, composed by families found in differential abundances in specific plastic substrates.

**Figure 5.**
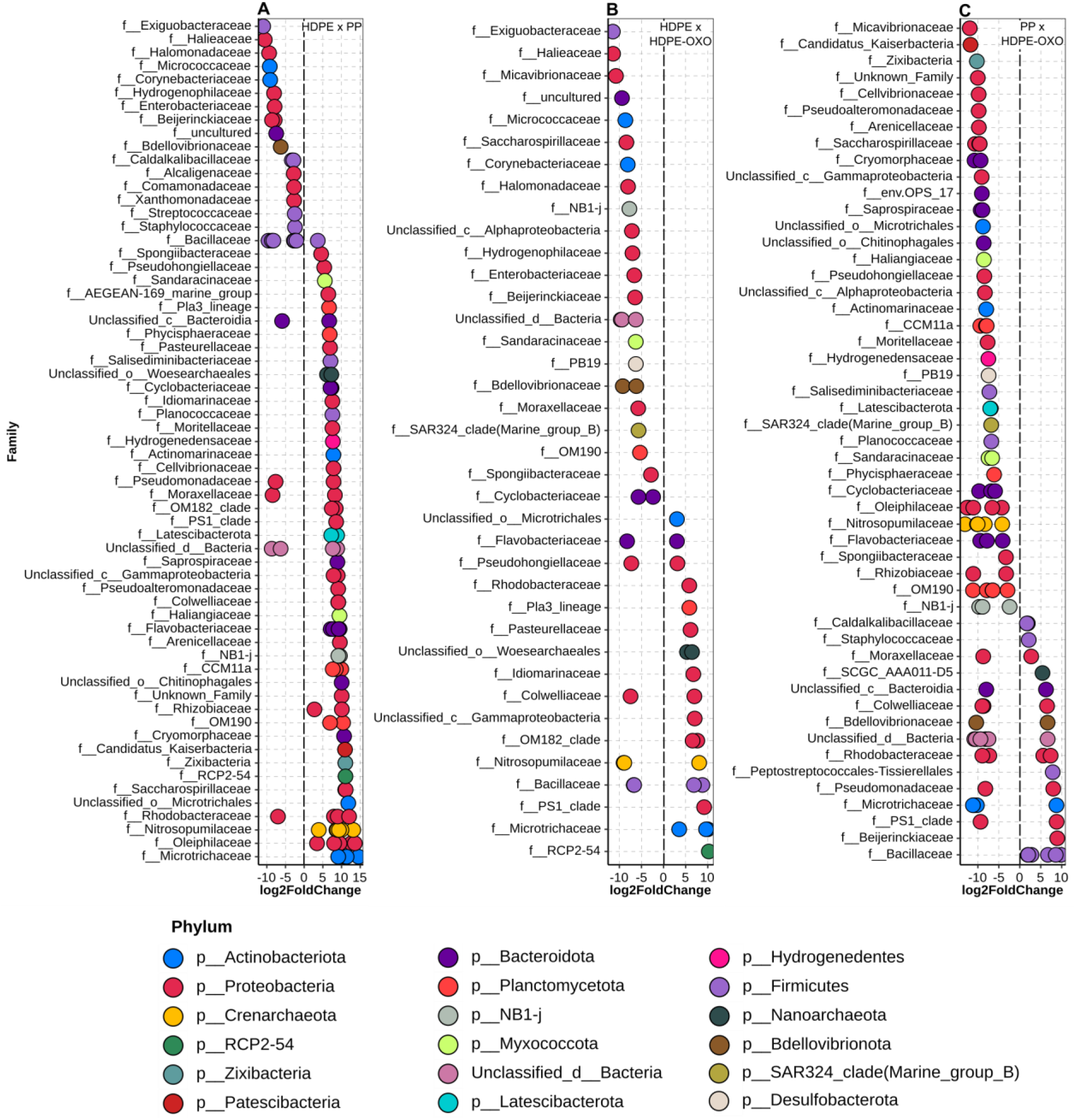
Differentially abundant Amplicon Sequence Variant (ASV) comparing HDPE and PP (A), HDPE and HDPE-OXO (B); PP and HDPE-OXO (C). Significant ASVs (padj < 0.05) are represented by single data points, grouped by family on the y-axis and by color according to the taxonomic phylum from which the ASV originates. Positive values (log2FoldChange) indicate ASVs that were significantly more abundant in HDPE (A and B) and PP (C) plastic substrates; Negative values indicate the opposite. Substrate: Gravel; AW, adjacent water; HDPE, High density polyethylene; HDPE-OXO, High density polyethylene with oxo-biodegradable additive BDA; PP, Polypropylene.

Fourteen families were classified into a generalist group, which included Bdellovibrionaceae, Halieaceae, Microtrichaceae, Pseudomonadaceae, and uncultured families. The HDPE plastic generalists were composed of 13 families, including CCM11a, Cryomorphaceae, Oleiphilaceae, Rhizobiaceae and Flavobacteriaceae, Nitrosopumilaceae. Among specialist families, the Beijerinckiaceae and Staphylococcaceae were significantly more abundant in PP samples, whereas NB1-j, OM190, Saccharospirillaceae, Spongiibacteraceae, Micavibrionaceae and unclassified class of Alphaproteobacteria were more abundant in HDPE-OXO samples. We did not identify families specifically associated with HDPE samples.

### 3.5. Core microbiome of the plastisphere community

To identify the core microbiome of the plastisphere from the deep Southwest Atlantic Ocean, the shared ASVs among plastic substrate types were examined (Figure 4). A total of 28 ASVs were shared among plastic types (Table S1), comprising the core microbiome of the plastisphere from the deep Southwest Atlantic Ocean. These ASVs were classified within 23 bacterial families, while 17 families included 97% of total core microbiome (Figure 4). This families were composed by Oleiphilaceae (30% of core microbiome), Rhizobiaceae (22%), unclassified Bacteria (13%), NB1-j (7%), Halieaceae (5%), Hyphomonadaceae (4%), Sneathiellaceae (3%), unclassified Bacteroidia class (3%), Micrococcaceae (2%), AB1 (1%), OM190 (1%), Thermaceae (1%), Paenibacillaceae (1%), Haliangiaceae (1%), Microtrichaceae (1%), Nannocystaceae (1%) and Eel-36e1D6 (1%). Six other families represented approximately 3% of the core microbiome (Figure 4).

### 3.6. Cultivable plastic-associated Bacteria

Bacterial strains were cultivated from all substrates, including all plastic types and gravel (Table S2). Overall, 15 strains were affiliated according to their 16S rRNA gene sequences to a bacterial genus. The most dominant family identified among the strains was Halomonadaceae, recovered from all substrates. This family was represented by two genera, *Salinicola* and *Halomonas*. Four families were isolated only from plastic substrates, Flavobacteriaceae (HDPE and HDPE-OXO), Pseudoalteromonadaceae (HDPE), Marinobacteraceae (HDPE-OXO) and Rhodobacteraceae (PP). In contrast, Micrococcaceae and Bacillaceae were isolated only from gravel. The comparison of the sequences of all isolates with the SILVA rRNA database confirmed the identities of the isolates. A phylogenetic tree was constructed with the bacterial sequences deposited in the database most closely related to our isolates (Figure S3). All the bacterial genera isolated in this study have been reported associated with or degrading hydrocarbon compounds (Table S2).

## 4. Discussion

Deep-sea environments are characterized by low temperatures, high pressure, absence of light and the consequent absence of photosynthetic primary production (Corinaldesi, 2015). Together with a general reduction of organic matter input, these extreme conditions promote high selective pressures on the microbial community. The input of anthropogenic substrates (i.e., sources of carbon) into the deep sea, such as plastic substrates, creates new habitats (or food sources) to be colonized by microorganisms. However, the composition of these substrata may select pelagic microorganisms with specific features that allow them to colonize and metabolize this carbon source (Dussud et al., 2018a; Dussud et al., 2018b).

Although the input of organic matter is suggested to be low, the pool of organic matter in deep waters could be enough to support high microbial richness (Luna et al., 2012), as indicated by our results. The highest richness on the gravel samples indicates that more microbial taxa were capable of attaching to the gravel in comparison with the plastic substrates, suggesting that plastics likely offer a strong selective pressure to colonization by the surrounding microbial community (McCormick et al., 2014). This is also supported by the lower microbial evenness observed in the plastic substrates (Figure S1C). However, statistical differences in alpha-diversity indices between plastic substrates and AW were not observed. Despite the high variability among samples and a limited number of replicates, these results indicate that the substrate effect on microbial communities is reflected by taxonomic composition rather than by diversity indexes. To the best of our knowledge, there is no previous study that has experimentally addressed the microbial colonization on plastic substrates in deep waters. Several previous studies have reported on surface waters (either from laboratory or field experiments) a lower richness and diversity of plastic substrates in comparison with the surrounding waters, suggesting that plastic substrates select for a specific and less diverse microbial community (McCormick et al., 2014; Ogonowski et al., 2018; Zettler et al., 2013). Nevertheless, deep water is generally oligotrophic, while surface waters have a constant input of organic matter from primary production from the phytoplanktonic community and could support high microbial diversity. These general oligotrophic conditions could also be attributed to the lack of significant differences in microbial richness and diversity among sites, which are exposed to the same depths and water mass. Abiotic parameters associated with water mass are suggested to be the main driver of the microbial community in the pelagic system. The similarity of these drivers observed among the sites could thus explain the lack of significant differences found in our study, as well as the high similarity among AW samples from different sites (Figure 2). Although at different sites, all samples were deployed at the same depth (3300 m), under similar temperatures (3-4 °C), salinities (34.6-35 psu), oxygen concentrations (above 5 mL L^−1^) and with similar low nutrient levels (oligotrophic) (Gonzalez-Silvera et al., 2004; Krueger et al., 2015).

Our results showing the strong effect of substrate type on microbial community structure (PERMANOVA, R^2^=0.78, *P* < 0.001) indicates niche partitioning of microbial communities among substrates (Dussud et al., 2018a). The substrate-dependence has been reported by studies regarding environmental and controlled conditions (Dussud et al., 2018a; Dussud et al., 2018b; Kirstein et al., 2019, 2018; McCormick et al., 2014; Zettler et al., 2013), showing evidence of the selective effect of plastic substrates. Under natural environmental conditions, substrate-dependence has also been reported in studies that randomly collected PMD (Plastic marine debris) (Didier et al., 2017; Dussud et al., 2018a; Ogonowski et al., 2018; Zettler et al., 2013) or that deployed plastic substrates (Kirstein et al., 2019), showing that the microbial community attached to a plastic substrate are distinct from free-living seawater communities or those attached to other hard substrata. Besides the substrate composition, the shape of the plastic particle could also act as a driver of the microbial diversity. For example, it is possible that the differences in the microbial structure and composition found between HDPE and PP substrates could also be attributed to shape differences in the plastics, not only to differences in composition. The HDPE substrate used in our study was in the form of a film, while the PP was a pellet. Because we didn’t have the same plastic type with multiple shapes available among our samples, we could not evaluate the isolated effect of plastic shapes on microbial diversity. Some recent studies have indicated that particle shape affects the microbial biofilm thickness (Wright et al., 2020), but if this reflects in changes in the microbial community composition is as yet unknown.

The microbial communities from gravel samples were more similar to the plastic substrates, while adjacent water samples were particularly distinct (Figure 2). These results were expected, as the plastic samples and gravel were deployed for 719 days; water samples represented only a single moment in time, while the plastic and gravel samples are the results of cell deposition and a dynamic succession over a long period. The AW samples are therefore a type of control, providing information on the microbial taxa present in the water column before retrieving the substrates. Corroborating our results, Oberbeckmann et al., (2016) demonstrated significant differences of multiple taxonomic groups when comparing plastic biofilm communities and the surrounding seawater communities; although the bacterial communities attached to PET bottles were distinct from the free-living seawater communities, the authors also found that PET-associated communities were similar to other types of particle-associated or glass-bound communities collected in the surrounding seawater. Those results confirm the ability of pelagic microorganisms to colonize a range of substrates without specificity (Dussud et al., 2018a).

At deeper taxonomic levels, the differences between the plastic and gravel samples were strongly evident. We identified 37 families significantly more abundant in plastic substrates than in gravel samples (Figure 2B), such as such as Methylologellaceae, Colwelliaceae, Pseudomonadaceae, Haliangiaceae, Micrococcaceae, Halieceaea, Oleiphilaceae, Rhizobiaceae, Microtrichaeaed, Flavobacteriaceae, Rhodobacteraceae and unclassified families of Alpha- and Gammaproteobacteria. Corroborating our results, previous studies have shown that microplastics were mainly colonized by Alpha- and Gammaproteobacteria, which were shown to act as primary colonizers, and Flavobacteria (Bacteroidetes), which appeared to act as secondary colonizers (Lee et al., 2008). Additionally, bacterial families classified as Flavobacteriaceae, Pirellulaceae, Rhodobacteraceae (Alphaproteobacteria) and Microtrichaceae (Acidimicrobiia) were identified as the most dominant families on microplastic (PE) biofilms exposed for 135 days to the marine environment at 12 m depth (Tu et al., 2020).

When comparing our microbial taxa from plastic substrates with previous studies related to the plastisphere in epipelagic ecosystems we found several families in common, such as Microtrichaceae, Rhizobiaceae, Halieaceae, Spongiibacteraceae, Rhodobacteraceae, Micavibrionaceae, Flavobacteriaceae, Halomonadaceae, Kangiellaceae, Hyphomonadaceae, Comamonadaceae, Oleiphilaceae and Bacillaceae (Amaral-Zettler et al., 2020; Feng et al., 2020; Pinto et al., 2019; Rogers et al., 2020). The family Oleiphilaceae comprises members that obligately utilize hydrocarbons through the alkane hydroxylase (*AlkB*) pathway (Golyshin et al., 2002); its detection in our plastic samples likely indicates their potential role in degrading plastic substrates in deep-sea ecosystems. Further, among the families described by these authors, Rhodospirillaceae members were not detected in our samples, which is in agreement with their photosynthetic capacity and thus their prevalence in epipelagic ecosystems. We detected a few families in our plastic samples which were not identified in these previous studies, such as Nitrosopumilaceae. Nitrosopumilaceae members are widely distributed in several deep-sea environments and have an important role as primary producers through ammonia oxidation (Zhong et al., 2020). Their presence in our plastic substrates likely reflects the high abundance of this family in these ecosystems, which might favor their attachment to a variety of substrates available for colonization.

Remarkable differences were observed when we grouped the microbial families by their distributions amongst plastic types as generalists (significantly abundant on all plastic types, HDPE, HDPE-OXO and PP), HDPE plastic generalists (significantly abundant in HDPE and HDPE-OXO), and specialists (significantly abundant in specific plastic substrates). Members of Bdellovibrionaceae, Halieaceae, Microtrichaceae and Pseudomonadaceae families were identified as generalists. The ability to colonize and potentially metabolize the carbon from plastic polymers of different substrates in deep environments, under oligotrophic conditions, confer advantages on these microbes in comparison to the entire microbial community. Some of these families have been previously described in association with different types of microplastics from several locations (e.g. Dussud et al., 2018a; Jiang et al., 2018; Tu et al., 2020). Tu et al., (2020) found a high abundance of Microtrichaceae members within biofilms of polyethylene microplastics from coastal seawater in the Yellow Sea, China, with increasing abundance according to longer exposure periods (135 days). Halieaceae members were detected in polyethylene, polypropylene and polystyrene microplastics from the Yangtze estuary (Jiang et al., 2018) and polyethylene microplastics from the Yellow Sea (Tu et al., 2020), both in China. This family is composed by marine bacteria that are capable of assimilating propylene through alkene monooxygenase genes (Suzuki et al., 2019). In addition, we observed microbial families associated specifically to HDPE samples (with and without biodegradable additives). Those microbes, in contrast to the generalist group, are suggested to be more adaptive to colonizing HDPE polymers, with weak or no influence from biodegradable additives. The influence of biodegradable additives was observed in the specialist taxa group (i.e. those microbes more adapted to a specific polymer type). The presence of additives in the polymer compositions may support microbial dynamics over time (Dussud et al., 2018b). Additionally, those additives could be an extra source of nutrients that may reflect in the multiplication of the different microorganisms. Similar results were reported by Dussud et al., (2018b) that suggested a strong effect of the polymer type on the bacterial community, because the composition of microbial biofilm on LDPE and LDPE-OXO (PE with pro-oxidant additives) was completely distinct, while AA-LDPE-OXO (artificially aged LDPE-OXO) and PHBV (poly(3-hydroxybutyrate-co-3-hydroxyvalerate) showed higher similarity, all under controlled conditions. As observed by these authors for shallow waters, we also observed an influence of the plastic types HDPE and HDPE-OXO on selecting specific microbial taxa in deep waters.

A total of 28 ASVs were identified as core microbiome members in the plastisphere. Defining a common core microbiome in the plastisphere across different studies may be difficult, because variations between experimental designs make it difficult to compare studies directly, as do the variety of study-specific approaches used to define the core (Didier et al., 2017; Tu et al., 2020; Zettler et al., 2013). However, some taxa observed in our study were widely described by previous studies, which provides evidence of common core members of the plastisphere from both surface and deep waters. ASVs from the bacterial families Oleiphilaceae and Hyphomonadaceae were found as members of the core microbiome in our plastic samples. These families have members known to degrade hydrocarbons (Golyshin et al., 2002; Ozaki et al., 2007) or are frequently associated to plastic substrates in the marine environment (Bryant et al., 2016; De Tender et al., 2017; Oberbeckmann et al., 2018; Pinto et al., 2019; Zettler et al., 2013).

In addition, taxa reported from plastic substrates, but not in the core microbiome were also identified. For instance, Microtrichaceae were reported as a dominant taxon on a PE surface during the early phase of biofilm formation (Tu et al., 2020) and Sneathiellaceae colonized plastic debris along a transect through the North Pacific Subtropical Gyre (Bryant et al., 2016). Notably, taxa not previously described from plastic substrates were also identified. Micrococcaceae is a well-documented bacterial family inhabiting deep-sea sediments (Chen et al., 2005, 2016; Sass et al., 2001); their members were already identified in sediments from the Southwest Indian Ridge at depths ranging from 1,662 to 4,000 m (Chen et al., 2016), in a hypersaline 3,500 m depth site in the Mediterranean Sea (Sass et al., 2001), and were isolated from an Antarctic lake and deep-sea sediments from the tropical West Pacific (Chen et al., 2005). *Allorhizobium-Neorhizobium-Pararhizobium-Rhizobium* spp. (Rhizobiaceae), a taxon reported as nitrogen-fixing (Franche et al., 2009), was an abundant member of the core microbiome. The family Rhizobiaceae is commonly involved in plant-microbe interactions and was reported recently in marine environments (Kimes et al., 2015). In deep-sea environments, the species of Rhizobiaceae *Georhizobium profundi* was isolated from sediment collected at 4,524 m depth (Cao et al., 2020), but its association with plastic substrata were only described in freshwater environments (Wang et al., 2020; Wen et al., 2020). Moreover, another taxa that comprised our core microbiome was NB1-j, an uncultivated bacterial family that was previously found in Japan Trench sediment at 6,292 m depth (Yanagibayashi et al., 1999), and in 800 to 1,450 m depth sediments heavily impacted by an oil spill in the northern Gulf of Mexico (Handam et al., 2018). Finally, taxa from AB1 family (previously assigned as unclassified Alphaproteobacteria) and Eel-36e1D6 (previously assigned as unclassified environmental clone groups), which also comprised the core microbiome, were reported in deep-sea hydrothermal fields, as well as in ferromanganese crusts (Nitahara et al., 2011). Overall, these results highlight a significant number of deep-sea taxonomic groups that were not described by previous studies inhabiting the plastic substrates but were found inhabiting our plastic substrates in the deep SW Atlantic Ocean.

We identified some taxa in the core microbiome that might be potentially related to plastic degradation, according to previous studies. For example, *Arthrobacter* spp. (Micrococcaceae) isolated from plastic waste in the Gulf of Mannar, India, was reported degrading high-density polyethylene (HDPE); after 30 days incubation, they had reduced the weight of the substrate by 12% (Balasubramanian et al., 2010). In addition, members of the Halieaceae family that have known capabilities of assimilating propylene through alkene monooxygenase genes (Suzuki et al., 2019), were described in plastic substrates from the Yangtze estuary (Jiang et al., 2018) and Yellow Sea (Tu et al., 2020), both in China. Members of the Paenibacillaceae family, such as *Paenibacillus* spp. have shown high potential to degrade LPDE and HDPE when in consortia with *Pseudomonas* spp., *Stenotrophomonas* spp. and *Bacillus* spp. (Skariyachan et al., 2017). Furthermore, we were able to isolate bacteria from our plastic substrates that comprise families and genera previously described as colonizing or degrading hydrocarbon substrates (Table S2). The isolation of these bacteria indicates that they were at least viable in deep-sea conditions and are important members to be explored in future studies to reveal their plastic-degradation capacity.

Information about microbial communities associated with the plastic substrata in the deep-ocean is scarce in published research studies (Krause et al., 2020; Woodall et al., 2018). Additionally, results from other experiments deploying samples for a long period in deep sea environments have not yet been published to date. Our pioneer study showed that several taxonomic groups previously described as plastic colonizers in surface waters seem to also colonize the plastic substrates in the deep sea. However, we also identified some groups in the plastisphere that are typically found inhabiting deep-sea sediments, such as NB1-j, Rhizobiaceae and Eel-36e1D6 members, most of them still poorly characterized and not yet cultivated. In addition, 13% of taxa in the core microbiome were not classified to any microorganism previously deposited in the reference databases, which might indicate sequencing artefacts or that we identified potential novel groups not yet described. Our study addresses the gap in the knowledge of microbial colonization in plastics deployed for a long period in the deep sea, highlighting the taxa potentially involved with plastic degradation processes. However, further studies are needed to better understand their role in plastic colonization and degradation in deep-sea ecosystems.

In summary, this study reported for the first time that deep-sea microbial communities of the Southwest Atlantic Ocean are involved in the colonization of plastic substrates. Furthermore, the microbial community colonizing the plastic surfaces were distinct and dependent on polymer type, but the site where the samples were deployed had no effect on the microbial community. Our results demonstrated a core microbiome exclusively composed of low abundance taxa; some members were not previously described as associated with plastic substrates, while other bacterial families had previously been described degrading plastics, but not in deep-sea environments. Additionally, we were able to cultivate and isolate some of these bacterial families from our plastic substrates. Our results indicate that a specific microbial community from the deep-sea can attach, colonize and potentially degrade plastic substrates. It is important to note that some of those members were reported degrading plastics in controlled conditions, and their ability to degrade the plastic compounds under deep-water conditions remains unknown without further investigation. We provide the first evidence of an unexplored microbial community inhabiting the deep-sea plastisphere, which may be used as a baseline to future studies about the functionality and the potential of degradation of these microbial communities living in oligotrophic conditions, in the absence of sunlight, under high pressure and low temperature.

## CRediT authorship contribution statement

**Luana Agostini:** Conceptualization, Methodology, Investigation, Data curation, Writing - original draft, review & editing. **Julio Cezar Fornazier Moreira:** Formal analysis, Data curation, Writing - original draft, review & editing, Visualization. **Amanda Gonçalves Bendia:** Formal analysis, Writing - original draft, review & editing, Visualization. **Maria Carolina Pezzo Kmit:** Writing - original draft, review & editing. **Linda Gwen Waters:** Conceptualization, Methodology, Investigation, Writing - review & editing. **Marina Ferreira Mourão Santana:** Conceptualization, Methodology, Investigation, Writing - review & editing. **Paulo Yukio Gomes Sumida:** Conceptualization, Methodology, Writing - review & editing, Project administration, Funding acquisition. **Alexander Turra:** Conceptualization, Methodology, Writing - review & editing, Project administration, Funding acquisition. **Vivian Helena Pellizari:** Conceptualization, Methodology, Writing - review & editing, Supervision.

## Acknowledgments

This work has been funded by the São Paulo Research Foundation (FAPESP), project BioSuOr grant #2011/50185-1. This study was financed in part by the Coordenação de Aperfeiçoamento de Pessoal de Nível Superior - Brazil (CAPES) - Finance Code 001. LA was funded by CAPES. The authors would like to thank the crew of R/V Alpha Crucis and R/V Alpha Delphini from IOUSP, and the crew of PRV Almirante Maximiano (H41) from Brazilian Navy for sampling and Rosa C. Gamba and LECOM’s research team for all scientific support and assistance in the lab. We also acknowledge the support of the National Council for Scientific and Technological Development (CNPq) for scholarship grants to AT (#309697/2015-8 and #310553/2019-9) and LW (No. 150159/2015-3).

## References

Amaral-Zettler, L.A., Zettler, E.R., Mincer, T.J., 2020. Ecology of the plastisphere. Nat. Rev. Microbiol. 18, 139–151. https://doi.org/10.1038/s41579-019-0308-0

Anderson, M.J., 2001. A new method for non-parametric multivariate analysis of variance. Austral Ecol. 32–46.

Bergmann, M., Klages, M., 2012. Increase of litter at the Arctic deep-sea observatory HAUSGARTEN. Mar. Pollut. Bull. 64, 2734–2741. https://doi.org/10.1016/j.marpolbul.2012.09.018

Bolyen, E., Rideout, J.R., Dillon, M.R., et al., 2019. Reproducible, interactive, scalable and extensible microbiome data science using QIIME 2. Nat. Biotechnol. 37, 852–857. https://doi.org/10.1038/s41587-019-0209-9

Bryant, J. A., Clemente, T.M., Viviani, D.A., Fong, A.A., Thomas, K.A., Kemp, P., Karl, D.M., White, A.E., DeLong, E.F., 2016. Diversity and Activity of Communities Inhabiting Plastic Debris in the North Pacific Gyre. mSystems 1, 1–19. https://doi.org/10.1128/msystems.00024-16

Callahan, B.J., McMurdie, P.J., Rosen, M.J., Han, A.W., Johnson, A.J.A., Holmes, S.P., 2016. DADA2: High-resolution sample inference from Illumina amplicon data. Nat. Methods 13, 581–583. https://doi.org/10.1038/nmeth.3869

Cao, J., Wei, Y., Lai, Q., Wu, Y., Deng, J., Li, J., Liu, R., Wang, L., Fang, J., 2020. Georhizobium profundi gen. Nov., sp. nov., a piezotolerant bacterium isolated from a deep-sea sediment sample of the New Britain Trench. Int. J. Syst. Evol. Microbiol. 70, 373–379. https://doi.org/10.1099/ijsem.0.003766

Caporaso, J.G., Lauber, C.L., Walters, W.A., Berg-lyons, D., Lozupone, C.A., Turnbaugh, P.J., Fierer, N., Knight, R., 2010. Global patterns of 16S rRNA diversity at a depth of millions of sequences per sample. Proceedings of the national academy of sciences, 108(Supplement 1), 4516–4522. https://doi.org/10.1073/pnas.1000080107

Carson, H.S., Nerheim, M.S., Carroll, K.A., Eriksen, M., 2013. The plastic-associated microorganisms of the North Pacific Gyre. Mar. Pollut. Bull. 75, 126–132. https://doi.org/10.1016/j.marpolbul.2013.07.054

Chen, M., Xiao, X., Wang, P., Zeng, X., Wang, F., 2005. Arthrobacter ardleyensis sp. nov., isolated from Antarctic lake sediment and deep-sea sediment. Arch. Microbiol. 183, 301–305. https://doi.org/10.1007/s00203-005-0772-y

Chen, P., Zhang, L., Guo, X., Dai, X., Liu, L., Xi, L., Wang, J., Song, L., Wang, Y., Zhu, Y., Huang, L., Huang, Y., 2016. Diversity, biogeography, and biodegradation potential of actinobacteria in the deep-sea sediments along the southwest Indian ridge. Front. Microbiol. 7, 1–17. https://doi.org/10.3389/fmicb.2016.01340

Chiba, S., Saito, H., Fletcher, R., Yogi, T., Kayo, M., Miyagi, S., Ogido, M., Fujikura, K., 2018. Human footprint in the abyss: 30 year records of deep-sea plastic debris. Mar. Policy 0–1. https://doi.org/10.1016/j.marpol.2018.03.022

Cole, M., Lindeque, P., Halsband, C., Galloway, T.S., 2011. Microplastics as contaminants in the marine environment: A review. Mar. Pollut. Bull. 62, 2588–2597. https://doi.org/10.1016/j.marpolbul.2011.09.025

Conway, J.R., Lex, A., Gehlenborg, N., 2017. UpSetR: An R package for the visualization of intersecting sets and their properties. Bioinformatics 33, 2938–2940. https://doi.org/10.1093/bioinformatics/btx364

Corinaldesi, C., 2015. New perspectives in benthic deep-sea microbial ecology. Front. Mar. Sci. 2, 1–12. https://doi.org/10.3389/fmars.2015.00017

Dang, H., and Lovell, C. R. (2016). Microbial surface colonization and biofilm development in marine environments. Microbiology and Molecular Biology Reviews, 80(1), 91–138.

De Tender, C.A., Devriese, L.I., Haegeman, A., Maes, S., Ruttink, T., Dawyndt, P., 2015. Bacterial Community Profiling of Plastic Litter in the Belgian Part of the North Sea. Environ. Sci. Technol. 49, 9629–9638. https://doi.org/10.1021/acs.est.5b01093

De Tender, C., Schlundt, C., Devriese, L.I., Mincer, T.J., Zettler, E.R., Amaral-Zettler, L.A., 2017. A review of microscopy and comparative molecular-based methods to characterize “Plastisphere” communities. Anal. Methods 9, 2132–2143. https://doi.org/10.1039/C7AY00260B

Didier, D., Anne, M., Alexandra, T.H., 2017. Plastics in the North Atlantic garbage patch: A boat-microbe for hitchhikers and plastic degraders. Sci. Total Environ. 599–600, 1222–1232. https://doi.org/10.1016/j.scitotenv.2017.05.059

Dussud, C., Meistertzheim, A.L., Conan, P., Pujo-Pay, M., George, M., Fabre, P., Coudane, J., Higgs, P., Elineau, A., Pedrotti, M.L., Gorsky, G., Ghiglione, J.F., 2018a. Evidence of niche partitioning among bacteria living on plastics, organic particles and surrounding seawaters. Environ. Pollut. 236, 807–816. https://doi.org/10.1016/j.envpol.2017.12.027

Dussud, Claire, Hudec, C., George, M., Fabre, P., Higgs, P., Bruzaud, S., Delort, A.M., Eyheraguibel, B., Meistertzheim, A.L., Jacquin, J., Cheng, J., Callac, N., Odobel, C., Rabouille, S., Ghiglione, J.F., 2018b. Colonization of non-biodegradable and biodegradable plastics by marine microorganisms. Front. Microbiol. 9, 1–13. https://doi.org/10.3389/fmicb.2018.01571

Eriksen, M., Lebreton, L.C.M., Carson, H.S., Thiel, M., Moore, C.J., Borerro, J.C., Galgani, F., Ryan, P.G., Reisser, J., 2014. Plastic Pollution in the World’s Oceans: More than 5 Trillion Plastic Pieces Weighing over 250,000 Tons Afloat at Sea. PLoS One 9, 1–15. https://doi.org/10.1371/journal.pone.0111913

Feng, L., He, L., Jiang, S., Chen, J., Zhou, C., Qian, Z.J., Hong, P., Sun, S., Li, C., 2020. Investigating the composition and distribution of microplastics surface biofilms in coral areas. Chemosphere 252, 126565. https://doi.org/10.1016/j.chemosphere.2020.126565

Franche, C., Lindström, K., Elmerich, C., 2009. Nitrogen-fixing bacteria associated with leguminous and non-leguminous plants. Plant Soil 321, 35–59. https://doi.org/10.1007/s11104-008-9833-8

Galgani, F., Souplet, A., Cadiou, Y., 1996. Accumulation of debris on the deep sea floor off the French Mediterranean coast. Mar. Ecol. Prog. Ser. 142, 225–234.

GESAMP, 2019. Guidelines or the monitoring and assessment of plastic litter and microplastics in the ocean (Kershaw P.J., Turra A. and Galgani F. editors). Rep. Stud. GESAMP 99, 130.

Golyshin, P.N., Chernikova, Ta.N., Abraham, W.R., Lünsdorf, H., Timmis, K.N., Yakimov, M.M., 2002. Oleiphilaceae fam. nov., to include Oleiphilus messinensis gen. nov., sp. nov., a novel marine bacterium that obligately utilizes hydrocarbons. Int. J. Syst. Evol. Microbiol. 52, 901–911. https://doi.org/10.1099/ijs.0.01890-0

Gonzalez-Silvera, A., Santamaria-Del-Angel, E., Garcia, V.M.T., Garcia, C.A.E., Millán-Nuñez, R., Muller-Karger, F., 2004. Biogeographical regions of the tropical and subtropical Atlantic Ocean off South America: Classification based on pigment (CZCS) and chlorophyll-a (SeaWiFS) variability. Cont. Shelf Res. 24, 983–1000. https://doi.org/10.1016/j.csr.2004.03.002

Hamdan, L.J., Salerno, J.L., Reed, A., Joye, S. B., Damour, M. The impact of the Deepwater Horizon blowout on historic shipwreck-associated sediment microbiomes in the northern Gulf of Mexico. Sci Rep 8, 9057 (2018). https://doi.org/10.1038/s41598-018-27350-z

Jambeck, J.R., Geyer, R., Wilcox, C., Siegler, T.R., Perryman, M., Andrady, A., Narayan, R., Law, K.L., 2015. Plastic waste inputs from land into the ocean. Science (80-.). 347, 768–771. https://doi.org/10.1126/science.1260352

Jiang, P., Zhao, S., Zhu, L., Li, D., 2018. Microplastic-associated bacterial assemblages in the intertidal zone of the Yangtze Estuary. Sci. Total Environ. 624, 48–54. https://doi.org/10.1016/j.scitotenv.2017.12.105

Kimes, N.E., López-Pérez, M., Flores-Félix, J.D., Ramírez-Bahena, M.H., Igual, J.M., Peix, A., Rodriguez-Valera, F., Velázquez, E., 2015. Pseudorhizobium pelagicum gen. nov., sp. nov. isolated from a pelagic Mediterranean zone. Syst. Appl. Microbiol. 38, 293–299. https://doi.org/10.1016/j.syapm.2015.05.003

Kirstein, I.V., Wichels, A., Gullans, E., Krohne, G., Gerdts, G., 2019. The plastisphere – Uncovering tightly attached plastic “specific” microorganisms. PLoS One 14, 1–17. https://doi.org/10.1371/journal.pone.0215859

Kirstein, I. V., Wichels, A., Krohne, G., Gerdts, G., 2018. Mature biofilm communities on synthetic polymers in seawater - Specific or general? Mar. Environ. Res. 147–154. https://doi.org/10.1016/j.marenvres.2018.09.028

Krause, S., Molari, M., Gorb, E. V., Gorb, S.N., Kossel, E., Haeckel, M., 2020. Persistence of plastic debris and its colonization by bacterial communities after two decades on the abyssal seafloor. Sci. Rep. 10, 1–15. https://doi.org/10.1038/s41598-020-66361-7

Krueger, M.C., Harms, H., Schlosser, D., 2015. Prospects for microbiological solutions to environmental pollution with plastics. Appl. Microbiol. Biotechnol. 99, 8857–8874. https://doi.org/10.1007/s00253-015-6879-4

Kumar, S., Stecher, G., Li, M., Knyaz, C., Tamura, K., 2018. MEGA X: Molecular evolutionary genetics analysis across computing platforms. Mol. Biol. Evol. 35, 1547–1549. https://doi.org/10.1093/molbev/msy096

Lee, J.W., Nam, J.H., Kim, Y.H., Lee, K.H., Lee, D.H., 2008. Bacterial communities in the initial stage of marine biofilm formation on artificial surfaces. J. Microbiol. 46, 174–182. https://doi.org/10.1007/s12275-008-0032-3

Love, M.I., Anders, S., Huber, W., 2014. Differential analysis of count data - the DESeq2 package, Genome Biology. https://doi.org/110.1186/s13059-014-0550-8

Luna, G.M., Bianchelli, S., Decembrini, F., De Domenico, E., Danovaro, R., Dell’Anno, A., 2012. The dark portion of the Mediterranean Sea is a bioreactor of organic matter cycling. Global Biogeochem. Cycles 26, 1–14. https://doi.org/10.1029/2011GB004168

McCormick, A., Hoellein, T.J., Mason, S.A., Schluep, J., Kelly, J.J., 2014. Microplastic is an abundant and distinct microbial habitat in an urban river. Environ. Sci. Technol. 48, 11863–11871. https://doi.org/10.1021/es503610r

Nitahara, S., Kato, S., Urabe, T., Usui, A., Yamagishi, A., 2011. Molecular characterization of the microbial community in hydrogenetic ferromanganese crusts of the Takuyo-Daigo Seamount, northwest Pacific. FEMS Microbiol. Lett. 321, 121–129. https://doi.org/10.1111/j.1574-6968.2011.02323.x

Oberbeckmann, S., Labrenz, M., 2020. Marine Microbial Assemblages on Microplastics: Diversity, Adaptation, and Role in Degradation. Ann. Rev. Mar. Sci. 12, 209–232. https://doi.org/10.1146/annurev-marine-010419-010633

Oberbeckmann, S., Kreikemeyer, B., Labrenz, M., et al., 2018. Interactions of microplastic debris throughout the marine ecosystem. Front. Microbiol. 1, 1–8. https://doi.org/10.1038/s41559-017-0116

Oberbeckmann, S., Kreikemeyer, B., Labrenz, M., 2017. Environmental factors support the formation of specific bacterial assemblages on microplastics. Front. Microbiol. 8, 2709. https://doi.org/10.3389/FMICB.2017.02709

Oberbeckmann, S., Osborn, A.M., Duhaime, M.B., 2016. Microbes on a Bottle : Substrate, Season and Geography Influence Community Composition of Microbes Colonizing Marine Plastic Debris. PLoS One 1–24. https://doi.org/10.1371/journal.pone.0159289

Ogonowski, M., Motiei, A., Ininbergs, K., Hell, E., Gerdes, Z., Udekwu, K.I., Bacsik, Z., Gorokhova, E., 2018. Evidence for selective bacterial community structuring on microplastics. Environ. Microbiol. https://doi.org/10.1111/1462-2920.14120

Ozaki, S., Kishimoto, N., Fujita, T., 2007. Change in the Predominant Bacteria in a Microbial Consortium Cultured on Media Containing Aromatic and Saturated Hydrocarbons as the Sole Carbon Source. Microbes Environ. 22, 128–135. https://doi.org/10.1264/jsme2.22.128

Pierdomenico, M., Casalbore, D., Chiocci, F.L., 2019. Massive benthic litter funnelled to deep sea by flash-flood generated hyperpycnal flows. Sci. Rep. 9, 1–10. https://doi.org/10.1038/s41598-019-41816-8

Pinnell, L.J., Turner, J.W., 2019. Shotgun metagenomics reveals the benthic microbial community response to plastic and bioplastic in a coastal marine environment. Front. Microbiol. 10. https://doi.org/10.3389/fmicb.2019.01252

Pinto, M., Langer, T.M., Hüffer, T., Hofmann, T., Herndl, G.J., 2019. The composition of bacterial communities associated with plastic biofilms differs between different polymers and stages of biofilm succession. PLoS One 14, 1–20. https://doi.org/10.1371/journal.pone.0217165

PlasticEurope, 2019. Plastics-the Facts 2019 – An analysis of European plastics production, demand and waste data.

Quero, G.M., Luna, G.M., 2017. Surfing and dining on the “plastisphere”: Microbial life on plastic marine debris. Adv. Oceanogr. Limnol. 8, 199–207. https://doi.org/10.4081/AIOL.2017.7211

Roager, L., Sonnenschein, E.C., 2019. Bacterial Candidates for Colonization and Degradation of Marine Plastic Debris. Environ. Sci. Technol. 53, 11636–11643. https://doi.org/10.1021/acs.est.9b02212

Roesch, L.F.W., Dobbler, P.T., Pylro, V.S., Kolaczkowski, B., Drew, J.C., Triplett, E.W., 2020. pime: A package for discovery of novel differences among microbial communities. Mol. Ecol. Resour. 20, 415–428. https://doi.org/10.1111/1755-0998.13116

Rogers, K.L., Carreres-Calabuig, J.A., Gorokhova, E., Posth, N.R., 2020. Micro-by-micro interactions: How microorganisms influence the fate of marine microplastics. Limnol. Oceanogr. Lett. 5, 18–36. https://doi.org/10.1002/lol2.10136

Sass, A. M., Sass, H., Coolen, M. J. L., Cypionka, H., Overmann, J. (2001). Microbial Communities in the Chemocline of a Hypersaline Deep-Sea Basin (Urania Basin, Mediterranean Sea). Applied and Environmental Microbiology, 67(12), 5392–5402. https://doi.org/10.1128/AEM.67.12.5392-5402.2001

Skariyachan, S., Setlur, A.S., Naik, S.Y., Naik, A. A., Usharani, M, Vasist, K. S. Enhanced biodegradation of low and high-density polyethylene by novel bacterial consortia formulated from plastic-contaminated cow dung under thermophilic conditions. Environ Sci Pollut Res 24, 8443–8457 (2017). https://doi.org/10.1007/s11356-017-8537-0

Schlining, K., von Thun, S., Kuhnz, L., Schlining, B., Lundsten, L., Jacobsen Stout, N., Chaney, L., Connor, J., 2013. Debris in the deep: Using a 22-year video annotation database to survey marine litter in Monterey Canyon, central California, USA. Deep. Res. Part I Oceanogr. Res. Pap. 79, 96–105. https://doi.org/10.1016/j.dsr.2013.05.006

Schmidt, C., Krauth, T., Wagner, S., 2017. Export of Plastic Debris by Rivers into the Sea. Environ. Sci. Technol. 51, 12246–12253. https://doi.org/10.1021/acs.est.7b02368

Sekiguchi, T., Sato, T., Enoki, M., Kanehiro, H., Kato, C., 2010. Procedure for isolation of the plastic degrading piezophilic bacteria from deep-sea environments. J. Japanese Soc. Extrem. 9, 25–30.

Shade, A., Handelsman, J., 2012. Beyond the Venn diagram: The hunt for a core microbiome. Environ. Microbiol. 14, 4–12. https://doi.org/10.1111/j.1462-2920.2011.02585.x

Suzuki, T., Yazawa, T., Morishita, N., Maruyama, A., Fuse, H., 2019. Genetic and physiological characteristics of a novel marine propylene-assimilating halieaceae bacterium isolated from seawater and the diversity of its alkene and epoxide metabolism genes. Microbes Environ. 34, 33–42. https://doi.org/10.1264/jsme2.ME18053

Tu, C., Chen, T., Zhou, Q., Liu, Y., Wei, J., Waniek, J.J., Luo, Y., 2020. Biofilm formation and its influences on the properties of microplastics as affected by exposure time and depth in the seawater. Sci. Total Environ. 734. https://doi.org/10.1016/j.scitotenv.2020.139237

Urbanek, A.K., Rymowicz, W., Mironczuk, A.M., 2018. Degradation of plastics and plastic-degrading bacteria in cold marine habitats. Appl. Microbiol. Biotechnol. 102, 7669–7678. https://doi.org/doi.org/10.1007/s00253-018-9195-y

Van Cauwenberghe, L., Vanreusel, A., Mees, J., Janssen, C.R., 2013. Microplastic pollution in deep-sea sediments. Environ. Pollut. 182, 495–499. https://doi.org/10.1016/j.envpol.2013.08.013

Wang, L., Luo, Z., Zhen, Z., Yan, Y., Yan, C., Ma, X., Sun, L., Wang, M., Zhou, X., Hu, A., 2020. Bacterial community colonization on tire microplastics in typical urban water environments and associated impacting factors. Environ. Pollut. 265, 114922. https://doi.org/10.1016/j.envpol.2020.114922

Wen, B., Liu, J.H., Zhang, Y., Zhang, H.R., Gao, J.Z., Chen, Z.Z., 2020. Community structure and functional diversity of the plastisphere in aquaculture waters: Does plastic color matter? Sci. Total Environ. 740, 140082. https://doi.org/10.1016/j.scitotenv.2020.140082

Woodall, L.C., Jungblut, A.D., Hopkins, K., Hall, A., Robinson, F., Gwinnett, C., Paterson, G.L.J., 2018. Deep-sea anthropogenic macrodebris harbours rich and diverse communities of bacteria and archaea. PLoS One 1–11. https://doi.org/10.1371/journal.pone.0206220

Woodall, L.C., Sanchez-vidal, A., Paterson, G.L.J., Coppock, R., Sleight, V., Calafat, A., Rogers, A.D., Narayanaswamy, B.E., Thompson, R.C., 2014. The deep sea is a major sink for microplastic debris. R. Soc. Open Sci. 1. https://dx.doi.org/10.1098/rsos.140317

Wright, R. J., Erni-Cassola, G., Zadjelovic, V., Latva, M., & Christie-Oleza, J. A. (2020). Marine Plastic Debris: A New Surface for Microbial Colonization. Environmental Science & Technology, 54(19), 11657–11672. https://doi.org/10.1021/acs.est.0c02305

Yanagibayashi, M., Nogi, Y., Li, L., Kato, C., 1999. Changes in the microbial community in Japan Trench sediment from a depth of 6292 m during cultivation without decompression. FEMS Microbiol. Lett. 170, 271–279. https://doi.org/10.1016/S0378-1097(98)00556-4

Zettler, E.R., Mincer, T.J., Amaral-Zettler, L.A., 2013. Life in the “ Plastisphere ” : Microbial Communities on Plastic Marine Debris. Environ. Sci. Technol. 47, 7137–7146. https://doi.org/dx.doi.org/10.1021/es401288x

Zhong, H., Lehtovirta-Morley, L., Liu, J., Zheng, Y., Lin, H., Song, D., Todd, J., Tian, J., Zhang, X.-H., 2020. Novel insights into the Thaumarchaeota in the deepest oceans: their metabolism and potential adaptation mechanisms 1–16. https://doi.org/10.21203/rs.3.rs-16227/v2

